# DNA aptamers for the recognition of HMGB1 from *Plasmodium falciparum*

**DOI:** 10.1101/528778

**Authors:** Diego F. Joseph, Jose A. Nakamoto, Oscar Andree Garcia Ruiz, Katherin Peñaranda, Ana Elena Sanchez-Castro, Pablo Soriano Castillo, Pohl Milón

## Abstract

Rapid Diagnostic Tests (RDTs) for malaria are restricted to a few biomarkers and antibody-mediated detection. However, the expression of commonly used biomarkers varies geographically and the sensibility of immunodetection can be affected by batch-to-batch differences or limited thermal stability. In this study we aimed to overcome these limitations by identifying a potential biomarker and by developing molecular sensors based on aptamer technology. Using gene expression databases, ribosome profiling analysis, and structural modeling, we find that the High Mobility Group Box 1 protein (HMGB1) of *Plasmodium falciparum* is highly expressed, structurally stable and steadily present along all blood-stages of *P. falciparum* infection. To develop biosensors, we used *in vitro* evolution techniques to produce DNA aptamers for the recombinantly expressed HMG-box, the conserved domain of HMGB1. An evolutionary approach for evaluating the dynamics of aptamer populations suggested three predominant aptamer motifs. Representatives of the aptamer families were tested for binding parameters to the HMG-box domain using microscale thermophoresis and rapid kinetics. Dissociation constants of the aptamers varied over two orders of magnitude between nano- and micromolar ranges while the aptamer-HMG-box interaction occurred in less than 30 seconds. The specificity of aptamer binding to the HMG-box of *P. falciparum* compared to its human homolog depended on pH conditions. Altogether, our study proposes HMGB1 as a potential biomarker and a set of sensing aptamers that can be further developed into rapid diagnostic tests for *P. falciparum* detection.

## Introduction

Malaria is an infectious disease that affects animals and humans, caused by protozoans of the genus *Plasmodium*. Malaria remains the cause of 435,000 deaths worldwide, with 219 million cases reported during 2017 [1]. Accurate treatment requires the identification of the parasite of the genus *Plasmodium*, in addition to the species causing the disease. Currently, there are several methods to diagnose malaria, such as PCR-based, Giemsa microscopy, and Rapid Diagnostic Tests (RDTs). Among these options, the last two appear to be suitable for low-income and mostly affected countries; nonetheless, they also present certain limitations [2].

Giemsa microscopy is inexpensive to perform, can differentiate malaria species and stages, and can quantify the parasites [2]. However, well-trained personnel is required and, frequently unavailable in low-income countries [2]. RDTs, on the other hand, detect *Plasmodium spp*. biomarker antigens in a small amount of blood using immobilized antibodies. The two most used biomarkers are histidine-rich protein 2 (HRP-2) and lactate dehydrogenase (LDH), which have been shown of high sensitivity and specificity for detecting *P. falciparum* [3]. However, *P. falciparum* strains lacking the HRP-2 and HRP-3 genes have appeared, increasing false negative results [4,5].

Additionally, the production of sensing antibodies requires advanced facilities to ensure both, reproducibility and consistency between batches. This limitation is aggravated by delivery logistics to the final user where transportation and temperature conditions affect the efficiency, sensibility, and specificity of RDTs. Although RDTs continue being the best alternative for rapid screening and detection of malaria in low-income countries, they require novel biomarkers for *Plasmodium spp*. to overcome the detection of false negatives and the development of sensing molecules that are more reproducible and thermostable.

Recent genomic, transcriptomic, and proteomic studies of *Plasmodium spp*. are a great source to identify potential new and abundant biomarkers [6–8]. In this context, ribosome profiling (RP) appears to fill the gap between transcriptomics and proteomics [9]. RP evaluates mRNA expression and estimates the ribosomal load of each particular mRNA [9]. Thus, RP allows the researcher to identify which genes are expressed at any given conditions and, at the same time, RP estimates the relative protein abundance potential. In the context of biomarker discovery, RP arrives as a valid alternative to identify proteins that are highly synthesized in the cell.

Aptamers offer a valid alternative for the development of biosensors as they overcome such immunogenic requirement of antibodies [10]. Additionally, aptamers are chemically synthetized which leads to the substantial reduction of variability between production batches [10]. Aptamers, identified through Systematic Evolution of Ligands by Exponential enrichment (SELEX) [11,12], have shown additional and significant advantages over antibodies without compromising binding and biochemical specificity parameters [10]. Aptamers for malaria detection have been successfully developed for the commonly used biomarkers HRP-2 and LDH [13–15]. Here, we use gene expression databases and recent RP reports to identify a potential new biomarker for the detection *Plasmodium spp*., recombinantly produce it to purity, and develop high-affinity aptamers as biosensors. Altogether, our study proposes a novel biomarker and a set of biosensors that can further be developed into RDTs for *Plasmodium spp*. detection.

## Material and Methods

### Biomarker selection

Translational gene expression regulation, as assessed by RP on asexual blood stages, were primarily used to identify highly translated mRNAs [8]. RNA-seq on all blood stages of *P. falciparum* were compared using PlasmoDB search tools [8,16,17]. Genes whose transcription was homogenous and constant among all blood stages of *P. falciparum* were selected. The corresponding proteins were probed for the availability of three-dimensional structures or the potential to be predicted. Proteins without atomic resolution or predicted structures were excluded as available structural reports show that aptamers bind to tertiary folds of proteins [18]. Protein stability and solubility, as well as the presence of transmembrane domains, were estimated using the ExPASy ProtParam tool (Swiss Institute of Bioinformatics, Switzerland) and TMHMM 2.0 (Center for Biological Sequence Analysis, Technical University of Denmark, DK). The human homolog of HMGB1 was used for negative selection steps. HMGB1 contains the HMG-box, which is a conserved domain among eukaryotes [19]. Both conserved domains, from *Plasmodium falciparum* or *Homo sapiens*, are subsequently going to be called HMG-box *Pf* and HMG-box *Hs*, respectively. Clustal Omega (EMBL-EBI, UK) was used to compare HMG-box *Pf* and HMG-box *Hs* by multiple sequence alignment analysis between these two domains [20].

### Oligonucleotides, cloning, expression, and purification of HMG-box domain

Oligonucleotides, unmodified or chemically modified with fluorescein, were synthesized by Macrogen (Korea). DNA sequences coding for HMG-box *Pf* and *Hs* were cloned into the plasmid pET-24c (+) (KanR) by GenScript (USA). The coding sequences of both domains were optimized for codon usage of the host (*E. coli* BL21) using OptimumGeneTM (GenScript, USA) and a His-Tag was added at the carboxyl end for further purification. *E. coli* BL21 (DE3) were made competent and transformed using Mix & Go *E. coli* Transformation Kit & Buffer Set (Zymo Research, USA). Transformed *E. coli* were cultured in Terrific Broth (yeast extract 24 g/L, tryptone 20 g/L, KH_2_PO_4_ 0.017 M, K_2_HPO_4_ 0.072 M) medium at 37°C until the absorbance at 600 nm reached 0.5. Then, Isopropyl β-D-1-thiogalactopyranoside (IPTG) 1 mM was added to induce protein expression and incubation continued for three hours. After incubation, the culture was centrifuged at 6,000 g for 20 min at 4 °C. Then, the supernatant was discarded, and the pellet was resuspended using HAKM_10_ Buffer (50 mM HEPES (pH 7.4), 70 mM ammonium acetate, 30 mM NaCl, 10 mM MgCl_2_, 6 mM 2-mercaptoethanol). Cells were pelleted again before lysis with using 5 mL BugBuster^®^ Master Mix (EMD Millipore, USA) for each gram of cells. After resuspension and incubation at room temperature for 20 minutes, the lysate was centrifuged at 12,000 g for 40 min at 4 °C. The supernatant was collected for purification using a 1 mL His Trap™ FF Crude Column containing Ni-NTA (GE Healthcare, USA). Briefly, the column was equilibrated using five volumes of His-Tag Buffer with 10 mM imidazole (20 mM sodium phosphate (pH 7.4), 0.5 M NaCl). Then, the collected supernatant was loaded into the column followed by three volumes of His-Tag Buffer containing 30 mM imidazole to remove nonspecific bound proteins. Finally, HMG-box was eluted with five volumes of His-Tag Buffer containing 300 mM imidazole. 1 mL fractions were collected independently and analyzed by SDS-PAGE electrophoresis (18% acrylamide) and Coomassie blue staining. Eluted proteins were dialyzed using a 3 kDa membrane D-Tube Dialyzer (Merck, Germany) to remove the imidazole in Storage Buffer (200 mM ammonium acetate, 50 mM HEPES, 6 mM 2-Mercaptoethanol, 10% glycerol, pH 7.1). Protein purity was assessed by (18% acrylamide) SDS-PAGE electrophoresis and Coomassie blue staining.

### Protein labeling

HMG-box *Pf* and *Hs* preparations were labeled with NHS-Biotin for SELEX rounds or with NHS-Atto540Q essentially as described in [21]. Protein preparations were dialyzed using NHS Labeling Buffer (200 mM NaCl, 50 mM HEPES, 6 mM 2-Mercaptoethanol, 5% Glycerol, pH 7.8) to remove primary amines. Labeling reactions were performed using a 2:1 label to protein ratio. Reactions were incubated with moderated stirring at room temperature for 1 hour. After incubation, non-protein-bound labels were quenched using Glycine in a final concentration of 25 mM. Labeled proteins were purified using His Trap™ FF Crude Column containing Ni-NTA (as described above) and dialyzed in Storage Buffer.

### SELEX

SELEX was performed using a 40-mer ssDNA random library (Trilink Biotechnologies, USA) which contained constant flanking regions: 5’ TAG GGA AGA GAA GGA CAT ATG AT (N40) TTG ACT AGT ACA TGA CCA CTT GA 3’. Amplification primers were: forward (5’ TAG GGA AGA GAA GGA CAT ATG AT 3’) and reverse (5’ TCA AGT GGT CAT GTA CTA GTC AA 3’). For transcription-mediated amplification of libraries, a T7 promoter sequence was added (T7 reverse primer: 5’ TTC AGG TAA TAC GAC TCA CTA TAG GGT CAA GTG GTC ATG TAC TAG TCA A 3’). Each selection round was composed of: i) Aptamer selection using biotinylated HGM-box that were immobilized on streptavidin-coated magnetic beads (Streptavidin Mag Sepharose™ from GE Healthcare, USA). The bead binding capacity for either HMG-box *Pf* or HMG-box *Hs* was estimated to19 pmol/μL beads slurry. Selections were performed in SELEX Buffer (100 mM NaCl, 1.5 mM KCl, 5 mM MgCl2, 6mM 2-Mercaptoethanol, 50 mM Tris HCl, pH 7.5) at 25°C with moderate and continuous shaking. ii) HGM-box-bound aptamers were separated from the solution using a magnet followed by resuspension using SELEX buffer and a second magnetic separation. The number of bead resuspensions (washes) varied along SELEX rounds (S2 Table). iii) Bound oligonucleotides were used for further regeneration of the ssDNA library by PCR using Trilink forward, and T7 Trilink-modified reverse primers using the Maxima Hot Start Green PCR Master Mix (2X) (Thermo Fisher Scientific, USA). iv) Resulting dsDNA were used for *in vitro* transcription using the TranscriptAid T7 High Yield Transcription Kit (Thermo Fisher Scientific, USA). v) *in vitro* reverse transcription was used to regenerate a ssDNA pool of enriched aptamers using the RevertAid Reverse transcriptase (Thermo Fisher Scientific, USA). Generally, the selection stringency was increased along cycles of SELEX by modifying ionic strength, time of incubation and the number of washes (S2 Table). Purity and size of dsDNA, RNA, and ssDNA was assessed by 8M UREA PAGE followed by methylene blue staining. Concentration was measured by absorbance at 260 nm in a Nanodrop One spectrophotometer (Thermo Fisher Scientific, USA).

### Diversity Visualization by Endonuclease (DiVE) assay and library enrichment

DiVE protocol was standardized based on Lim *et al*. [22]. All the experiments were performed using the Thermo Scientific S1 endonuclease (Thermo Fisher Scientific, USA). For the reaction tubes, 200 ng of double-stranded DNA (dsDNA) from each SELEX cycle was used. The final volume of the reaction was 20 μL, and each sample contained S1 Reaction Buffer (40 mM sodium acetate (pH 4.5), 300 mM NaCl, 2 mM ZnSO_4_). Each tube reaction was subjected to denaturation at 95°C for 5 minutes and reannealing at 65°C for 5 minutes. After that, 1 μL of S1 endonuclease U/μL was added to the mixture followed by 30 minutes incubation at 65°C. Reactions were stopped by adding ethylene-diaminetetraacetic acid (EDTA) to a final concentration of 2 mM (pH 8.0). Finally, 5μL of each sample was mixed with 1 μL of RunSafe (Cleaver Scientific, UK) and visualized in a 2% agarose gel. Low diversity samples appeared as defined bands of the expected DNA length while high diversity samples appeared as smears or were absent.

### Library sequencing

Cycles four, six, eight, ten, and fourteen were taken for Next Generation Sequencing as they showed low (cycle four), partial (cycles six and eight) and high (cycles ten and fourteen) enrichment as evaluated by DiVE. The dsDNA pools were processed as described in the 16S Metagenomic Sequencing Library Preparation protocol (Illumina, USA). Briefly, 5 ng of dsDNA from each selected cycle was amplified by PCR for eight cycles using forward (5’ TCG TCG GCA GCG TCA GAT GTG TAT AAG AGA CAG TAG GGA AGA GAA GGA CAT ATG AT 3’) and reverse (5’ GTC TCG TGG GCT CGG AGA TGT GTA TAA GAG ACA GTC AAG TGG TCA TGT ACT AGT CAA 3’) Trilink primers modified with overhang adapters. After amplification, PCR products were purified with the AMPure XP beads (Beckman Coulter, USA) protocol. Subsequently, sequencing indexes were added by PCR, provided by the Nextera^®^ (R) XT Index Kit (Illumina, USA). Finally, indexed PCR products were purified with AMPure XP beads (Ampure, USA). Each indexed PCR product was quantified by absorbance at 260 nm and diluted to a final concentration of 4 nM using 10 mM Tris (pH 8.5). Five microliters of each indexed PCR were mixed to obtain the pooled library. Finally, the pooled library was denatured using an equal volume of 0.2 N NaOH, incubated and room temperature for 5 minutes and diluted again to a final concentration of 4 pM using HT1 Buffer (Illumina, USA) and loaded into the MiSeq Reagent Nano Kit v2 cartridge (Illumina, USA) for 300 reading cycles.

### Bioinformatic analysis and aptamer selection

For a primary analysis of the sequences and a comparison between SELEX cycles, the bioinformatic toolkit FASTAptamer was employed [23]. Reverse data retrieved by sequencing was reverse complemented and mixed with the forward data to obtain a collection of sequences comprising both sequencing steps. The FASTAptamer Count command allowed to estimate the frequency for all sequences in each SELEX cycle. Subsequently, the FASTAptamer Enrich command was used to estimate the relative increase in abundance between subsequent SELEX cycles for each sequence analyzed. Candidate molecules were ranked depending on the enrichment between cycles and abundance. The MEME Suite and Clustal Omega (EMBL-EBI) was used to evaluate the presence of conserved motifs [20,24]. Mfold was used to determine the Gibbs free energy of potential secondary structures using the selection conditions, 25°C, 100 mM Na+, and 3 mM Mg+^2^ [25]. Conserved flanking regions of the aptamer library were included in the analysis.

### Three-dimensional structure modeling and docking predictions

Modeling of three-dimensional (3D) structures of aptamers followed the principles described by Jeddi *et al*. [26]. Briefly, the Mfold web server was employed to predict secondary structures of the aptamers as described above. Output files were saved using the Vienna bracket format [25]. A 3D equivalent single-stranded RNA (ssRNA) model was built for each aptamer through the RNAComposer web server, using the bracket structures previously obtained [27]. The 3D ssRNA models were manually transformed into ssDNA structures using the molefacture and the autoPSF plugin of VMD software [28]. This was achieved by replacing the H5 atom in each uracil residue with a methyl moiety, manually replacing the uracil denomination with thymine in the PDB file, and changing the ribose to deoxyribose with the autoPSF builder [26]. Then, we used VMD and NAMD to refine the new 3D DNA structures in an ionized (100 mM NaCl) water box through 10,000 iterations of energy minimization [29,30]. The 3D structure of HMG-box *Pf* was built using homology modeling with the SWISS-MODEL Suit ([31]. Finally, the ZDOCK server was employed to predict the structural interaction between the aptamers and the HMG-box *Pf* [32]. We used the ZDOCK server with ZDOCK 3.0.2 without specifying specific interactions.

### Determination of dissociation constants

Dissociation constants were measured using Microscale Thermophoresis (MST) on a Monolith NT.115 instrument (NanoTemper, Germany). Thermal migrations were monitored in real time by using a FAM-labeled probe (probe^FAM^) annealed to the 5’ end of the aptamer. Pre-annealing of the probe to the aptamer was performed by mixing aptamers with probe^FAM^ in 4:1 ratio followed by 5 min denaturation at 95°C and 20 min renaturation at room temperature. Reactions (20 μL) were prepared containing 50 nM of aptamers and varying concentrations of either HMG-box *Pf* or *Hs*. Before measuring, samples were centrifuged at 14,000 g for 5 min. The reactions were loaded into MO-K022 standard capillaries (NanoTemper, Germany) and measured. A preliminary fluorescence scan of capillaries was used to assess homogeneity within the titration. Reactions were excluded if a fluorescence difference between scans was higher than 10%. Time courses of fluorescence measurements were set 5 seconds with the infrared (IR) laser off, 30 seconds with the IR on, and 5 seconds off. A delay of 25 seconds between reads was used for thermal equilibration of the instrument. All reactions were performed in PBS at room temperature. IR laser power was set to 20% while fluorescence excitation power was set to 20% to 70% according to the dynamic of each aptamer.

### Kinetic Experiments

To measure the kinetics of aptamer-protein interactions an SX20 stopped-flow apparatus was used (Applied Photophysics, UK). All reactions were performed in SELEX Buffer (pH 7.4). Aptamers, PfR6 and PfE3, were pre-annealed to the probe^FAM^ as described above and called from herein, PfR6^FAM^ and PfE3^FAM^. Fluorescent measurements were carried out at 22 °C using a monochromatic light source (470 nm) powered with 10 mA. Emitted fluorescence was measured with photomultipliers set at 320 V after passing a long-pass filter with a cut-off of 515 nm essentially as described in [33]. Equal volumes (60 μL) of each reactant were rapidly mixed and fluorescence changes over time were measured. Typically, one solution contained aptamer (0.25 μM), probe^FAM^ (0.2 μM) while the second solution was composed of 1 μM HMG-box^540Q^ *Pf* or *Hs*, quenching derivatives of their respective proteins labeled with Atto-540Q. Between 7 to 10 replicates were measured. Every single measurement acquired 1,000 data points in a logarithmic sampling mode. Apparent rate constants were estimated by non-linear regression with exponential equations using Prism 7.02 (Graphpad Sofware, USA). Averaged rate constants were calculated as described in [34].

## Results

### HMGB1 as a biomarker for *Plasmodium* detection

Recent advancements in the ‘omics field allow novel approaches to assess gene expression regulation at several stages, namely, gene transcription, mRNA translation and protein availability in the cell. Among these, translation regulation can be studied by RP, which provides unprecedented parameters, both qualitative and quantitative [35]. At any given cell growth state, RP can relatively estimate the ribosome load for each identified mRNA [35]. Recent applications of RP for *P. falciparum* identified more than 3,000 actively translated mRNAs at different stages of parasite life-cycle [8]. Actively translated mRNAs shall result in highly expressed proteins, an appealing feature to propose new potential biomarkers as parasite density can be estimated. Using the freely available RP databases, biomarker candidates were ranked according to their expression in the merozoite blood-stage, as being the stage with the most active protein synthesis apparatus (Fig 1A). Any gene whose expression was null during any blood stage was excluded since homogenous expression during all blood stages was desired. To assess the expression variability along blood-stages of *P. falciparum* the most expressed genes from RP datasets were validated by transcriptomic reports using PlasmoDB [16,17]. Candidates whose expression was not uniform during all blood stages were eliminated. Biomarker candidates that met both selection criteria, being abundant and showing similar expression patterns during all blood stages of the parasite, were pre-selected (Fig 1).

**Fig 1.**
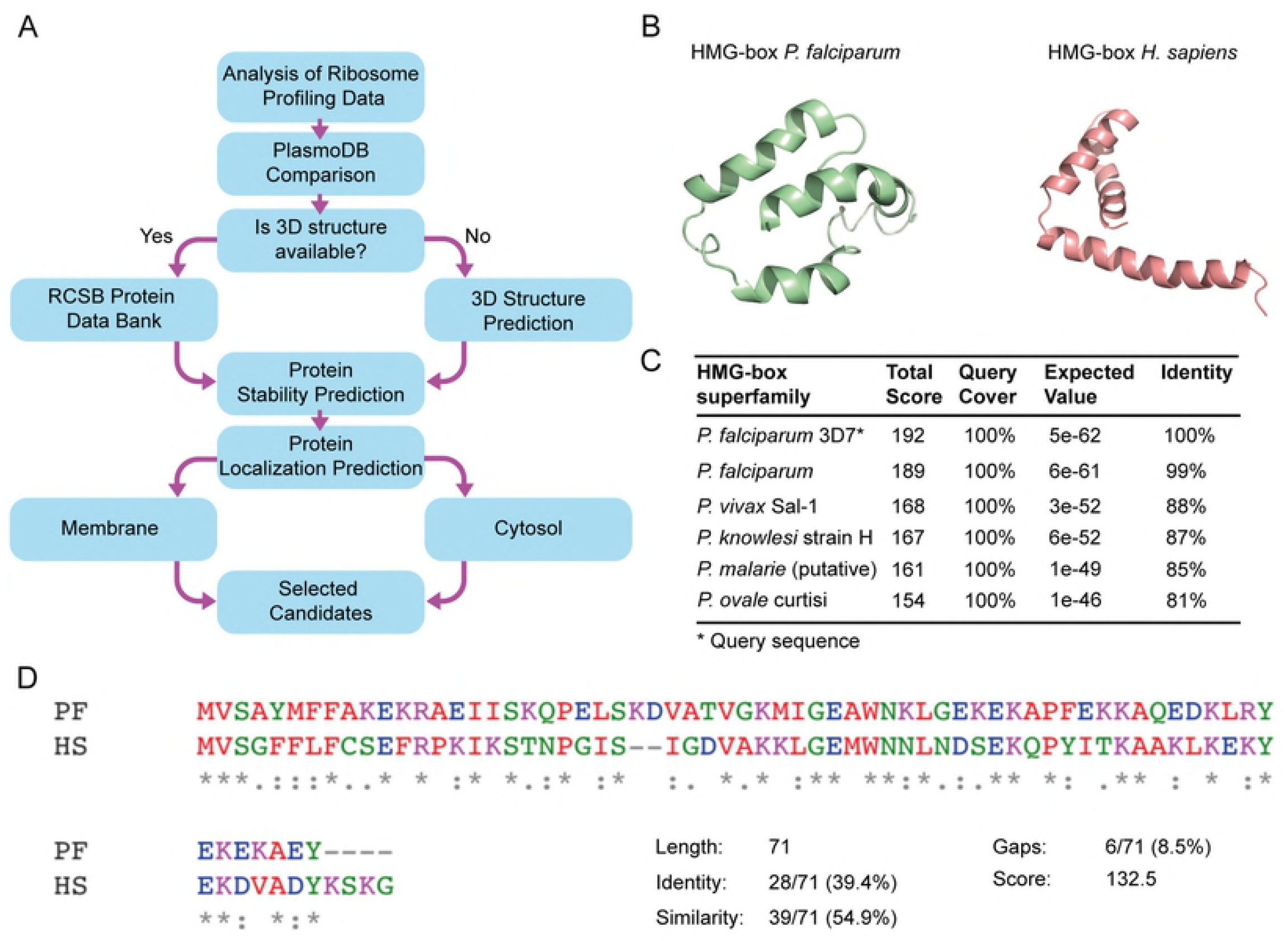
HMG-box as a potential biomarker of *Plasmodium falciparum*. (A) Workflow for the identification of highly abundant proteins of *Plasmodium falciparum* as novel biomarker candidates. (B) Structure predictions by homology of HMG-box from *P. falciparum* and its human homolog using the Swiss-model suite [31]. (C) BLASTp suite of HMG-box *Pf* shows a high percentage of identity with other pathogenic *Plasmodium* species. (D) Clustal Omega (EMBL-EBI, UK) multiple sequence alignment for HMG-box *Pf* and HMG-box *Hs*. Both domains have 71 amino acids and share 39.4 *%* of identity and 54.9% of similarity.

Further bioinformatic analysis allowed to investigate structural features of the potential biomarkers. From the top five biomarker candidates, only fructose-bisphosphate aldolase (FBPA) had a known 3D structure (PDB 1A5C) [36]. However, the tertiary structure of HMGB1, eIF1, Tetraubiquitin, 14-3-3 protein was available from modeling based on proteins from other species that shared at least 40% of their amino acid sequence (Plasmo DB, [16]). From the five potential biomarkers of *P. falciparum*, HMGB1 showed up as being abundant, steady along all blood-stages of the parasite and, folding into a stable tertiary structure (Fig 1B). Therefore, HMGB1 displays substantial premises for the development of molecular sensors for *Plasmodium* detection. The HMGB1 protein from *Plasmodium spp*. has a human homolog, and both contain a DNA binding domain, HMG-box (Fig 1B) [19]. Furthermore, the HMG-box domain is highly conserved among *Plasmodium* species that infect humans, such as *P. vivax* (88% identity), *P. knowlesi* (87% identity), *P. malarie* (85% identity), and *P. ovale* (81% identity) (Fig 1C). Of particular importance, HMG-box *Pf* shows limited identity (39.4 %) with HMG-box *Hs* (Fig 1D). Altogether, abundance, availability during all infectious stages of the parasite, and low identity with the human counterpart make the HMG-box an appealing biomarker to develop new biosensors.

### Aptamers against HMG-box *Pf*

In order to select molecular sensors that can specifically recognize the HMG-box *Pf*, we used Systematic Evolution of Ligands by Exponential Enrichment (SELEX) combined with next-generation sequencing (NGS). SELEX allows to *in vitro* select single-stranded oligonucleotides that strongly bind a given molecule [12]. These oligonucleotides are known as aptamers [37] and were shown to have significant advantages over antibodies such as better thermal stability and synthesis consistency between production batches without compromising their binding affinities [10,38]. These characteristics are very appealing for further development of aptamers into rapid diagnostic tests to be deployed in rural areas in low-income countries, the most affected by Malaria. We performed fourteen SELEX cycles (Fig 2A) starting from a library of ssDNA containing 40 randomized nucleotides. In order to enhance the selection of tight binders, HMG-box *Pf* concentration, incubation time and strength of washes varied, increasing the selection stringency along the SELEX cycles (S2 Table). Whereas, to increase the specificity of the potential aptamers, negative selections using the human counterpart, HMG-box *Hs*, were introduced at cycles five, eight, ten, and fourteen.

**Fig 2.**
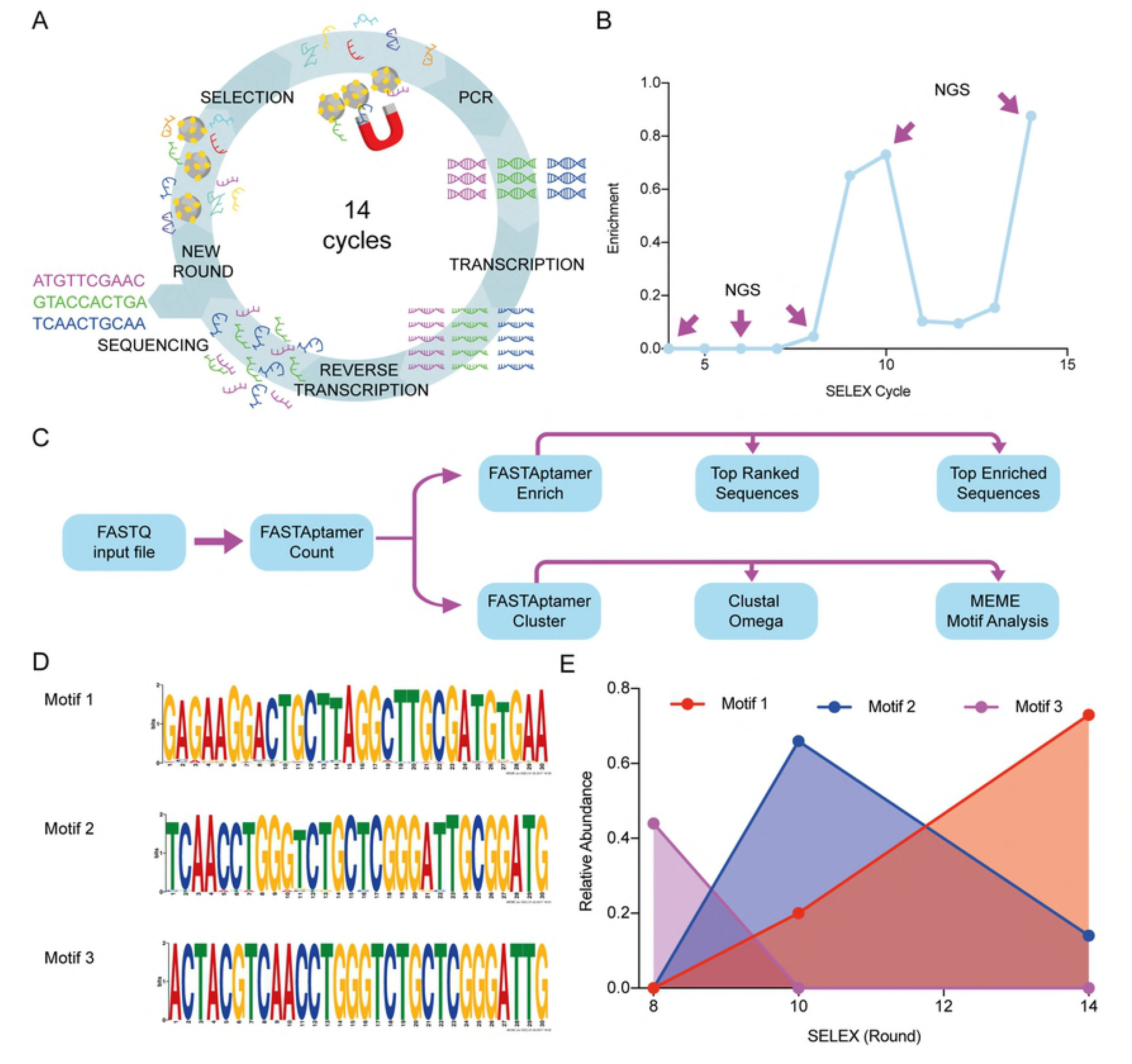
NGS allows the identification of aptamer motifs and their evolution during SELEX. (A) Workflow of the SELEX technique and variations applied in this work. The ssDNA library (N40) was incubated with the HMG-box proteins immobilized in magnetic beads. After incubation, a magnet is used to separate the ssDNA molecules that bound the proteins on the magnetic beads. To amplify the bound molecules a PCR using a modified reverse primer with the T7 promoter region is performed. After purification, *in vitro* transcription is carried out, followed by *in vitro* reverse transcription. Finally, an enriched ssDNA library is generated and used for a next cycle or for sequencing. (B) Progression of enrichment along steps of selection as evaluated by the DiVE technique [22]. The ratio between non-degraded DNA (enriched)/total DNA was calculated from image quantifications of the resulting agarose gel using the ImageJ software [39] (see material and methods). Rounds 4, 6, 8, 10 and 14 were chosen for NGS analysis. (C) Scheme depicting the workflow for sequence analysis. (D) MEME (The MEME Suite) was used to identify the most characteristic motifs in each sequenced SELEX cycle. Three motifs were the most represented in cycles 8 (motif 3), 10 (motif 2), and 14 (motif 1). Sequences from cycles 8 and 10 share a conserved region inside motifs 3 and 2, whereas this region is lost in motif 1. No representative motif was detected in cycle 4. (E) Dynamics of aptamer motifs along SELEX cycles.

It is expected that the initial variability of the starting library decreases along the SELEX cycles, enriching a sub-population that contains the potential aptamers. To assess the enrichment of aptamers over the high variability of the starting library, we used the DiVE assay [22] and bioinformatic analysis of sequencing data (see below). DiVE uses the S1 endonuclease to differentiate between homoduplex and heteroduplex dsDNA by cleaving short single-stranded ends, nicks, and heteroduplex loops. An enriched pool would have more dsDNA homoduplex compared with a non-enriched pool. Thus, the homoduplex would not be cleaved and could be visualized in an agarose gel. The DiVE assay allowed to confirm enrichment at cycles eight, ten, and fourteen with a loss of homogeneity from cycle eleven to thirteen (Fig 2B and S1 Fig.). The dsDNA obtained from cycles four, six, eight, ten, and fourteen were used for NGS using the Illumina system.

Raw sequences (typically over 80,000 reads for each SELEX cycle) obtained from sequencing were processed using the FASTAptamer toolkit (Fig 2C) [23]. Briefly, a FASTQ file was processed using the algorithm FASTAptamer Count that ranks DNA sequences based on their abundance for each SELEX cycle (Refer to S1 Data for raw ranked sequences). Best ranked sequences in cycle 14 varied between 14,995 RPM (Reads per Million, a normalized index for frequency) and 27,6943 RPM. In cycle ten, the most abundant sequences ranged between 5,933 RPM and 103,658 RPM. Final rounds of selection provided fewer sequences with higher frequency, while earlier selection cycles did not show sequences over 190 RPM of representation (For raw data, refer to S2 Data). An important indicator of aptamer selection can be surmised from the enrichment of each sequence along the selection procedure. This can be estimated from the ratio of sequence frequencies belonging to successive SELEX cycles. Thus, we used the FASTAptamer Enrich algorithm to calculate the enrichment of all aptamer sequences over two SELEX cycles, i.e., fourteen over ten or ten over eight. Most enriched sequences showed a frequency ratio ranging from 82 to 566 in cycles fourteen over ten. The enrichment ratio for cycles ten over eight showed increased enrichment ranging between 24.1 and 16.5 (See S3 Data for all sequences). Combining the results from both, ranking and enrichment, nine potential aptamers were identified (Table 1).

**Table 1.**
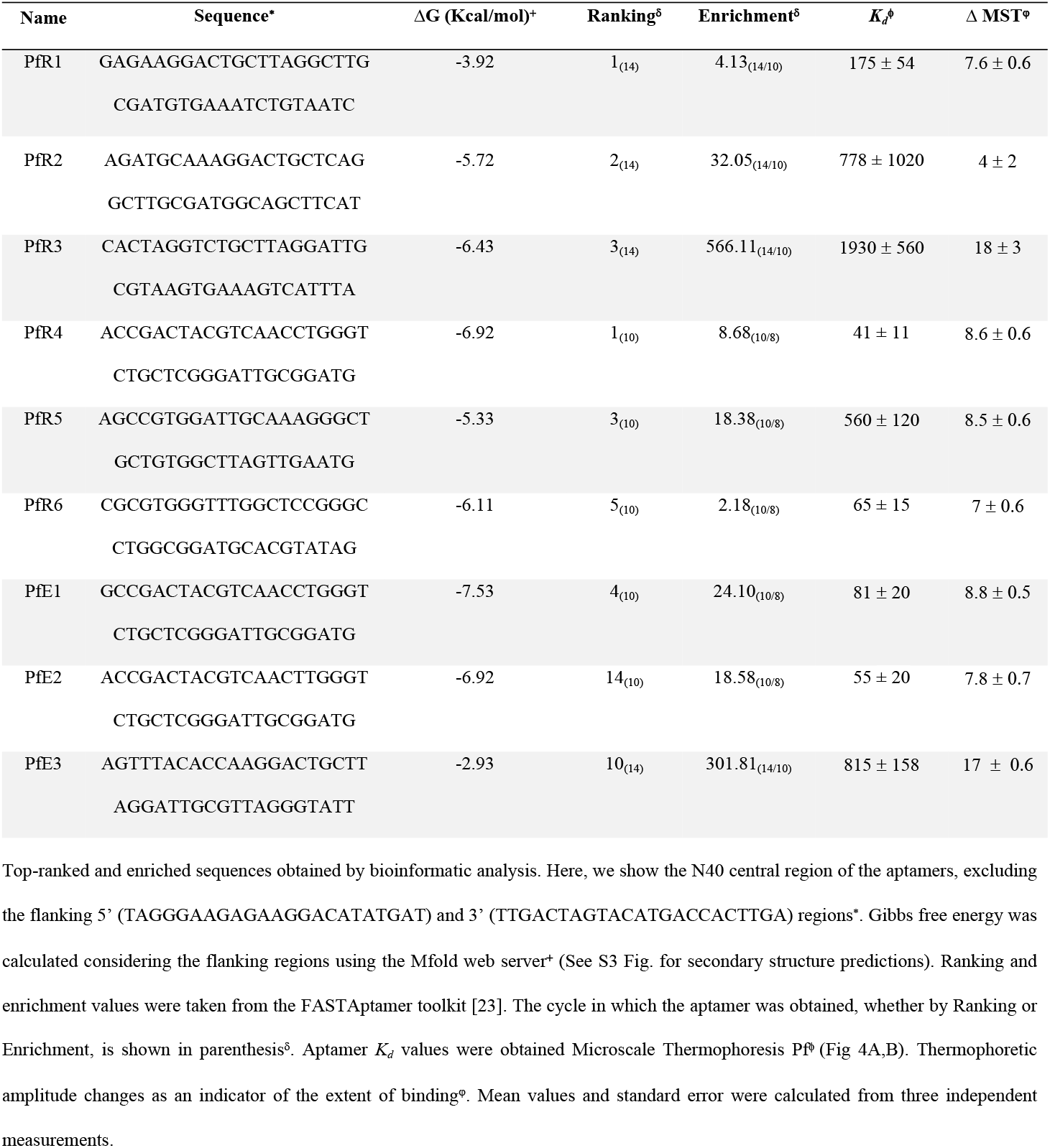
Top-ranked and enriched sequences as evaluated by NGS.

Additionally, sequences from the sequential selection process were also used to select for families of aptamers sharing a common and conserved sequence with few substitutions within the sequence. The variability within each family can arise either from the original library or by synthesis errors of the Taq polymerase, T7 RNA polymerase, or retro transcriptase that were used to regenerate the pool of ssDNA. To further analyze how families are enriched along the SELEX cycles of selection, we used the FASTAptamer Cluster algorithm [23] together with MEME Suite Motif Analysis and the Clustal Omega [20,24]. Representative motifs in cycles eight, ten, and fourteen were identified (Fig 2D). Motif 3 was the most abundant in cycle eight with about 40% of share. However, motif 3 diminished in cycles ten and fourteen, concomitantly with an increase of share for motif 2. A relative abundance of 70 % for this family was followed by a reduction to less than 20 % in cycle fourteen. On the other hand, motif 1, with low representation in cycles eight and ten, appeared highly abundant in cycle fourteen, with 80 % of share (Fig 2E). Motif 1 was represented by sequences PfR1, PfR2, PfR3, and PfE3. Whereas motif 2 was represented by sequences PfR4, PfR5, PfR6, PfE1, and PfE2. PfR4 and PfE1 shared motif 2 and 3 within their sequences (Table 1). Overall, the bioinformatic flow used here allowed to identify nine potential aptamers for the HMG-box *Pf*, with three representing families of aptamers that vary their relative abundances along the SELEX process (Table 1, Fig 2E).

### Interaction parameters between aptamers and HMG-box *Pf*

Nine aptamers, among the best, ranked or enriched, and belonging to either of the three most represented motifs were chemically synthesized and tested for binding to the HMG-box *Pf* using Microscale Thermophoresis (MST). MST allows the analysis of molecular interactions by measuring drifts of fluorescent reporters that are induced by small temperature changes, named thermophoresis [40,41]. The degree of thermophoresis change depends on the bound and unbound state of the fluorescent reporter with the ligand. Therefore, the thermophoresis changes as a function of ligand concentration allow calculating dissociation constants (*K*_d_) [40]. Several MST applications for aptamer research have been reported (see [42] for detailed methods). Here, we label the aptamer with a fluorescent probe (probe^FAM^) and use it at a constant concentration and increased the concentrations of HMG-box *Pf*, ranging from 0.1 nM to 2 μM. The measured affinities ranged from 50 nM to 2 μM, with aptamers PfR1, PfR4, PfR6, PfE1, and PfE2 being in the low nanomolar range and PfR2, PfR5, and PfE3 in the micromolar range of dissociation constants (Table 1) (Fig 3A,B). PfR6 was representative of the group showing high affinities, while aptamer PfE3 showed the greatest MST amplitude at a lower affinity.

**Fig 3.**
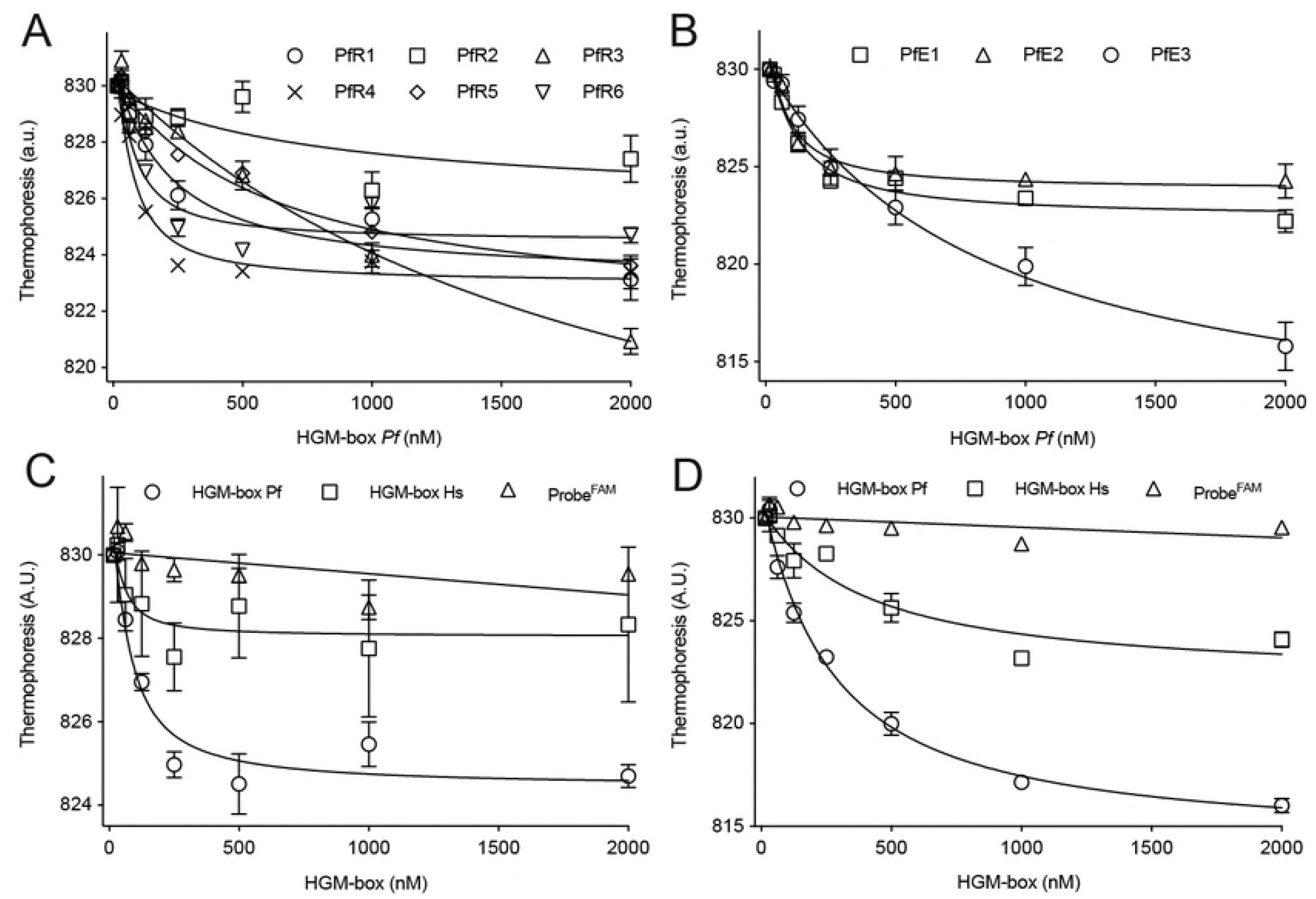
Aptamers efficiently bind the HMG-box from *P. falciparum*. MST analysis of the interaction between aptamers and HMG-box. (A) Thermophoresis changes of best ranked aptamers as a function of HMG-box *Pf* concentration. (B) Thermophoresis changes of the most enriched aptamers as a function of HMG-box *Pf* concentration. Specificity assessment for PfR6 (C) and PfE3 (D). The binding of each aptamer to HMG-box *Pf* (open circles) was compared to the human homolog (open squares) or in the absence of the aptamer (open triangles). All reactions were measured in triplicates. Error bars indicate standard deviations. Continuous lines show the non-linear regression using a quadratic equation, allowing to estimate a *K_d_* values (Table 1).

Another parameter to consider for optimal aptamers lays on specificity determinants. Thus, we probed PfR6 and PfE3 binding to HMG-box *Hs* by MST. The extent of binding PfR6 to the human homolog of HMG-box was lower when compared to the *Plasmodium* counterpart (Fig 3C). However, the PfE3 aptamer showed residual capacity to interact with HGM-box *Hs*, however with a 10-fold reduced affinity (Fig 3D). In brief, the PfR6 aptamer appears to strongly interact with the HMG-box from *P. falciparum*.

In order to further analyze the binding determinants of the PfR6 interaction with HMG-box *Pf*, we studied the kinetics of the reaction by the Stopped-flow (SF) technique. SF allows monitoring changes of fluorescence or absorbance in time and upon rapid mixing of two reactants. Here we used Föster Resonance Energy Transfer (FRET) between HMG-box (*Pf* or *Hs*) modified with the acceptor dye Atto540Q (from herein, HMG-box^540Q^ *Pf* or *Hs*) and the PfR6 aptamer pre-annealed with a small complementary probe labeled with Fluorescein (PfR6^FAM^) (see material and methods). The interaction between the fluorescent aptamer and the quenching HMG-box resulted in a fluorescence decrease in time, due to the quenching effect induced by the vicinity of the non-emitting acceptor Atto540Q to the fluorescein reporter in the aptamer. To assess the specificity of the signal, first, we mixed HMG-box^540Q^ *Pf* with the fluorescent probe only (in the absence of aptamer PfR6) and resulted in a negligible fluorescence change in time, indicating low or no interaction between the reactants (Fig 4A). Consistently, the mixing of PfR6^FAM^ with the unlabeled HMG-box *Pf* did not result in any fluorescence change in time. Mixing HMG-box^540Q^ *Pf* with PfR6^FAM^ resulted in a decrease of fluorescence over time, consistent with a rapid binding of PfR6^FAM^ with HMG-box^540Q^ *Pf*. Nonlinear analysis with an exponential function allowed estimating the apparent average rate of the interaction, *k_AV_* = 0.58 ± 0.01 s^−1^.

**Fig 4.**
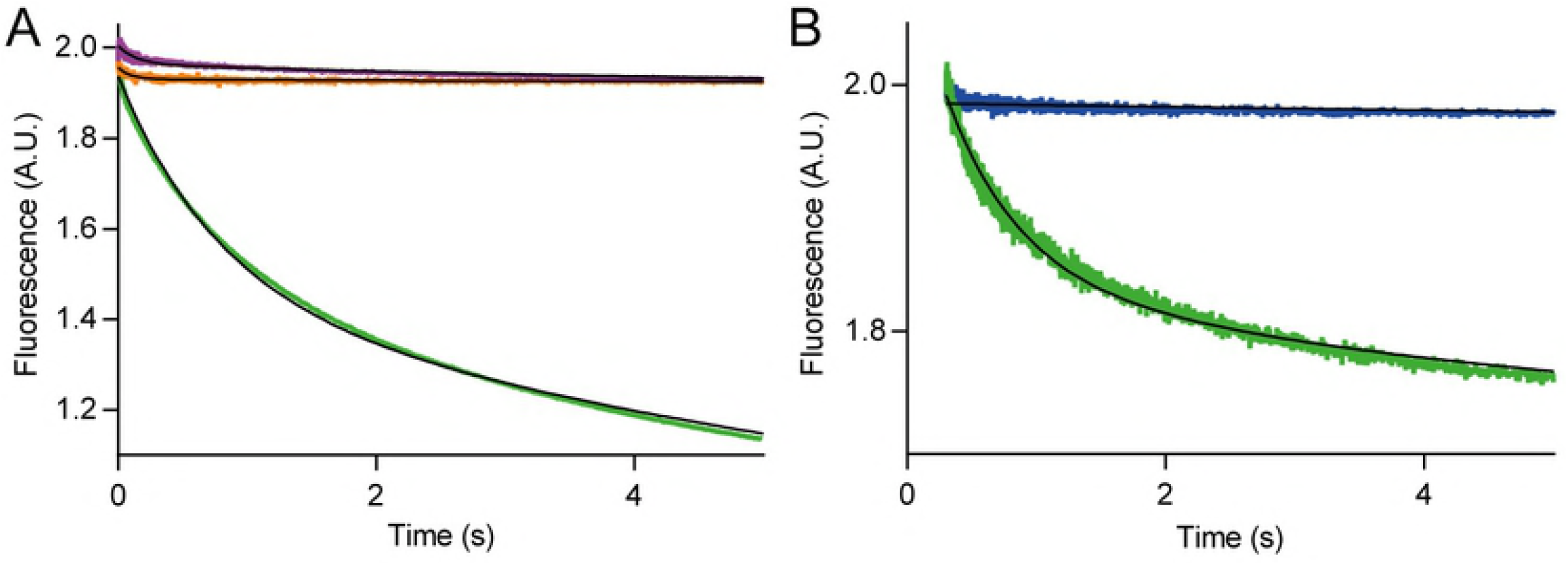
Kinetics of PfR6 interaction with HMG-box. HMG-box^540Q^ and PfR6^FAM^ were used to monitor the interaction between the aptamer and the protein in real-time using a stopped-flow apparatus (Applied Photophysics, UK). (A) FRET between HMG-box^540Q^ and PfR6^FAM^. Mixing the labeled protein with the probe-bound aptamer (green) resulted in a decrease of fluorescence over time, indicating that the binding of both components results in the quenching of fluorescein due to its vicinity with the quencher in the protein. FRET assessment controls lacking the aptamer or the quencher Atto-540 in the HMG-box are shown in orange and purple traces respectively. (B) HMG-box ^540Q^ *Pf* (green) or HMG-box ^540Q^ *Hs* (blue) comparison during the binding with PfR6^FAM^ at pH 5.5. 7-10 replicates for each reaction were measured and averaged. Continuous lines show the fitting with exponential functions.

Similar experiments using the human counterpart of the biomarker, HMG-box^540Q^ *Hs*, showed variable degrees of binding as a function of pH. At pH=5.5, PfR6 did not show any binding to HMG-box^540Q^ *Hs* while the binding to the plasmodium counterpart was maintained (Fig 4B). Thus, PfR6 appears to bind specifically to HMG-box^540Q^ *Pf* at the given conditions. However, the specificity of the binding was found to be perturbed by increasing the pH (S2 Fig.). Nonlinear regression showed an average rate constant *k*_AV_ = 0.18 ± 0.02 s^−1^ for PfR6 interaction with HMG-box^540Q^ *Pf*. This rate increased with pH, reaching *k_AV_* = 1.12 s^−1^. Ion concentrations and pH have been shown to influence the tertiary structure of aptamers, therefore affecting their binding properties [43]. Altogether, our results indicate that PfR6 interacts efficient- and rapidly with the HMG-box *Pf* domain and provide experimental conditions where the specificity of the interaction is greatly enhanced.

### Structural model of the PfR6–HMG-box *Pf* complex

In order to evaluate the theoretical molecular interactions between aptamers and HMG-box *Pf*, 3D models of the aptamers were generated followed by a molecular docking approximation. Secondary structures of the aptamers were predicted by MFold (S3 Fig.) [25]. Then, the RNAComposer tool was used to generate 3D RNA models of the aptamers. The resulting models were converted to DNA structures with VMD and then stabilized through 10,000 iterations of energy minimization as described by Jeddi I. and Saiz L. [26]. 2D and 3D predictions indicated that all nine aptamers showed unstructured regions at the 5’ end. The length of the unstructured regions varied between 10 and 26 bases (S3 and S5 Fig.). On the other hand, the 3’ end contained highly structured domains that included the 3’ flanking regions of the aptamer library (S5 Fig.). PfR6 showed 21 (out of 86) unstructured nucleotides at the 5’ end. The structured 3’ region showed three segments with short double-stranded helixes and several helix-to-helix interactions (Fig 5A). Thus, PfR6 appears to fold into significantly structured regions, at the same time, provides a long unstructured segment that could be exploited to further develop diagnostic tests.

**Figure 5.**
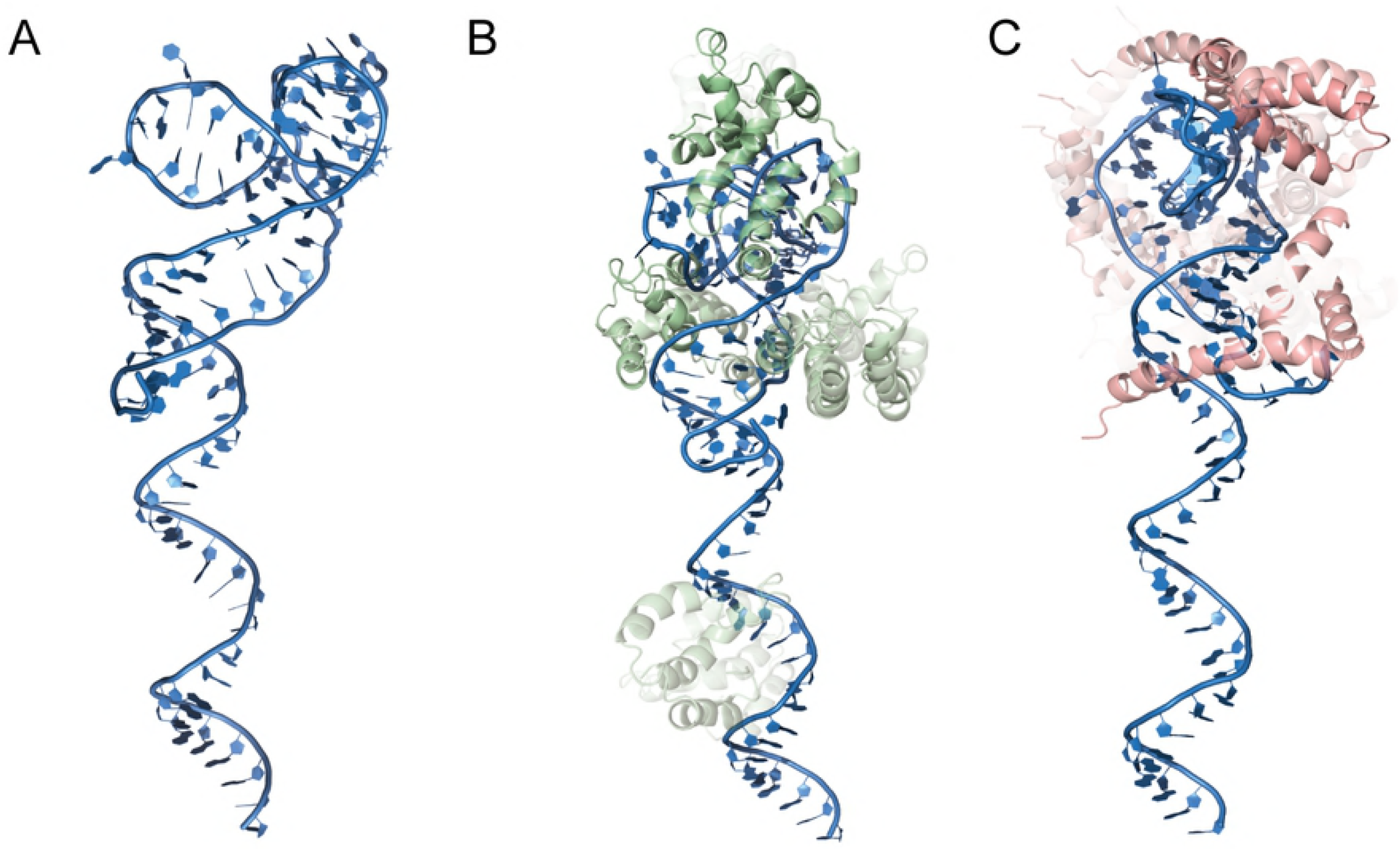
Structural modeling of aptamers and the PfR6–HMG-box *Pf* complex. (A) Tertiary structure prediction of the PfR6 aptamer. (B) Alternative models for the PfR6– HMG-box *Pf* complex as evaluated by ZDOCK [32]. The degree of opacity is proportional to the ZDOCK Score (10% opacity corresponding to a score of 1350, maximum opacity corresponds to a score of 1850) (S6 Fig.). (C) as B for the PfR6–HMG-box *Hs*. The PfR6 aptamer is depicted in sky blue, the HMG-box *Pf* in pale green, and HMG-box *Hs* in pink.

Modeling of the PfR6–HMG-box *Pf* complex was performed using the predicted tertiary structures of the aptamer and the protein (see above) using the ZDOCK tool [32]. We generated 2,000 complexes for each aptamer and evaluated their respective docking score based on rigid bodies interactions (S4 Data, S6 Fig.). The average value of the docking scores did not indicate substantial differences between the selected aptamers. However, the highest score varied for each aptamer, ranging from 1500 to 1850, with PfE3 and PfR6 among the highest ones (S6 Fig.). Thus, *in silico* docking results appear to be consistent with the biochemical data as measured by MST (Fig 3, Table 1). A comparison among the best ten PfR6–HMG-box *Pf* complexes, as evaluated by the ZDOCK score, indicates that in eight out ten models HMG-box *Pf* interacts with the structured region of PfR6 (Fig 5B). As a comparison, the PfR6–HMG-box *Hs* complex (human counterpart) showed much lower ZDOCK scores; however, the first ten models show an interaction of the protein with the structured region of the aptamer (Fig 5C).

## Discussion

The identification of potential biomarkers is a complex process where many factors must be considered. In the case of *Plasmodium* species, an ideal biomarker should establish the presence or absence of infection, determine the species involved, be detectable at low concentrations, and be proportional to parasite density [44]. To account for these characteristics, the successful entrepreneurship requires several experimental stages, from basic research towards final clinical validations. Initial steps for biomarker selection can be pursued using bioinformatic strategies, with large databases of the -omics providing valid alternatives [45,46]. Genome-wide translation dynamics studies as revealed by RP, and in contrast to transcriptomic approaches, offer a precise approximation for the *P. falciparum* proteome [8]. Secondary analysis of RP datasets allowed selecting HGMB1 as a highly expressed protein during all the blood stages of *P. falciparum* (Fig 1, S1 Table). Structural stability, conservation among *Plasmodium spp*., and diversity to human homologs provided additional properties to cope for an ideal biomarker (Fig 1). Conserved regions of proteins from *Plasmodium* could be exploited to generate biosensors for the genus. Conversely, non-conserved regions allow generating species-specific biosensors [47]. Alternative approaches for the selection of biomarkers for malaria have been described [48], leading to different candidates for biomarkers, however, considering biomarker abundance shall increase the detection likelihood of the corresponding biosensors. As a comparison, during merozoite blood-stage, HGMB1 showed 88.45 folds more expression than the commonly used PfLDH biomarker, while HRP-2 expression is null.

HMGB1 is a highly conserved non-histone protein among eukaryotes that binds the minor groove of DNA and enhances the activity of transcription factors [49]. HMGB1 of *H. sapiens* can be exported to the extracellular medium and can fulfill a role as an alarmin by being involved in infectious and non-infectious inflammatory conditions [50]. *In vivo* assays showed that HMGB1 *Pf* to be an inducing factor for potent pro-inflammatory processes, leading to the induction of TNF-α and Nitric Oxide Synthase [49]. Both, transcriptional enhancer and extracellular activities, add additional features to HMGB1 as a biomarker. One can expect that *Plasmodium spp*. exploit the natural regulatory network of the human HMGB1 by exporting HMGB1 *Pf*, although, to our knowledge, experimental evidence is still lacking. Deletion of the HMGB2 homolog from *Plasmodium* resulted in attenuated pathogenicity in mouse models [51], indicating an involvement on severe inflammatory responses. Altogether, HMGB1 and HMGB2 appear to be involved in the inflammatory response of the host, making the HMGB family a candidate for directed inhibitory drugs. Indeed, several aptamers are currently being evaluated in clinical trials [52]. Here, we showed that PfR6 tightly bind the conserved domain HMG-box displaying affinities that are comparable to antibodies used as biological drugs. Besides their potential for pharmacological administration, PfR6 may further be modified as a biosensor of HMGB1 from *P. falciparum*.

Aptamers for the detection of infectious diseases are of remarkably flexibility for the development of diagnostic assays. Such versatility arises from uncountable chemical modifications that are commercially available. Aptamer-tethered enzymes have been shown of great sensibility and to differentiate *P. falciparum* from *P. vivax* in blood samples [53]. Additionally, the development of aptamers using species specific epitopes allowed to isolate an aptamer that could accurately discriminate between the above species [47]. Aptamers against PfLDH were used in impedance spectroscopy setups allowing to propose electrochemical sensors that recognize the biomarker at a wide range of conditions [54]. Measurements of intramolecular FRET between aptamer linked dyes allowed the development of aptamer beacons [55]. In this work we show that fluorescently modified aptamers PfR6 and PfE3 allowed to monitor the binding to HGM-box *Pf* using two different techniques, providing substantial premises to develop aptamer beacons. Indeed, stopped-flow assays resulted in large FRET changes that, in principle, are compatible with further developing of detection assays. Additionally, PfR6 interaction with HMG-box *Pf* is rapid, taking place in less than a minute. Both, the resulting high signal and rapid binding, can allow the development RDTs with lower detection times.

The chemical flexibility of aptamers can furthermore be exploited for binding optimization and stability in complex samples such as blood, plasma, saliva, etc. Indeed, the introduction of unnatural modifications can result in increased diversity, higher affinity as well as providing inhibitory capabilities [56]. The introduction of a single phosphorodithioate substitution resulted in a 1000-fold increase of affinity for an RNA aptamer against thrombin, leading to picomolar dissociation constants [57]. PfR6 showed binding affinities in the nanomolar range which could be further increased by targeted substitutions.

RDTs are the best alternative for rapid screening and detection of malaria in emerging countries, and the present research allows to cope with two current limitations. First, HMG-box *Pf*, is proposed as an alternative biomarker to the mostly available HRP and LDH. Second, the development of aptamers and their fluorescent derivatives as potentially new biosensors of HMG-box *Pf*. The latter can be further investigated in order to develop alternative detection methods, i.e. enzyme-tethered, surface immobilized or molecular beacons.

## Author contributions

OAGR and PM conceived the project. DFJ, AESC, KP, OAGR, and PSC performed experiments. DFJ, JAN, KP, and PM analyzed the data. PM and DFJ wrote the manuscript with the inputs from all authors.

## Acknowledgments

We are very thankful to Andrey Konevega for kindly providing access to the Microscale Thermophoresis technology.

## Funding

This work was funded by grants from Grand Challenges Canada (0692-01-10), Fondecyt (038-2014-Fondecyt and 136-2016-Fondecyt), and InnóvatePerú (297-INNOVATEPERU-EC-2016) to PM.

## Competing Interests

BDM is a Startup of the Laboratory of Applied Biophysics and Biochemistry (UPC) where PM owns shares. BDM is funded by the Peruvian government (136-2016-Fondecyt) and is committed to develop diagnostic methods for neglected diseases.

## List of supporting information

S1 Fig. Aptamer enrichment analysis by the DiVE assay.

S2 Fig. Kinetics of PfR6 interaction with HMG-box Pf and Hs at pH 7.5.

S3 Fig. Secondary structures of top-ranked and enriched aptamers.

S4 Fig. MST analysis of the aptamer-HGM-box complex formation.

S5 Fig. Tertiary structures of top-ranked and enriched aptamers.

S6 Fig. Averaged ZDOCK scores for the PfR6–HMG-box *Pf* complex.

S1 Data. Aptamer frequency analysis by the Fastaptamer count algorithm.

S2 Data. Aptamer ranking for all sequenced SELEX round.

S3 Data. Aptamer enrichment analysis by the Fastaptamer Enrich algorithm.

S4 Data. ZDOCK scores for 2,000 iterations corresponding to the nine aptamers.

S1 Table. Potential biomarkers for *Plasmodium falciparum* infection as proposed by the algorithm in Fig 1.

S2 Table. Reaction conditions used in each SELEX cycle.

## References

1. World Health Organization. World malaria report 2018. 2018 Nov pp. 1–210.

2. Wongsrichanalai C, Barcus MJ, Muth S, Sutamihardja A, Wernsdorfer WH. A review of malaria diagnostic tools: microscopy and rapid diagnostic test (RDT). Am J Trop Med Hyg. 2007;77: 119–127.

3. Li B, Sun Z, Li X, Li X, Wang H, Chen W, et al. Performance of pfHRP2 versus pLDH antigen rapid diagnostic tests for the detection of Plasmodium falciparum: a systematic review and meta-analysis. Arch Med Sci. 2017;13: 541–549. doi:10.5114/aoms.2017.67279

4. Cheng Q, Gatton ML, Barnwell J, Chiodini P, McCarthy J, Bell D, et al. Plasmodium falciparum parasites lacking histidine-rich protein 2 and 3: a review and recommendations for accurate reporting. Malar J. BioMed Central; 2014;13: 283. doi:10.1186/1475-2875-13-283

5. Menegon M, L’Episcopia M, Nurahmed AM, Talha AA, Nour BYM, Severini C. Identification of Plasmodium falciparum isolates lacking histidine-rich protein 2 and 3 in Eritrea. Infect Genet Evol. 2017;55: 131–134. doi:10.1016/j.meegid.2017.09.004

6. Le Roch KG, Johnson JR, Florens L, Zhou Y, Santrosyan A, Grainger M, et al. Global analysis of transcript and protein levels across the Plasmodium falciparum life cycle. Genome Res. 2004;14: 2308–2318. doi:10.1101/gr.2523904

7. Bozdech Z, Mok S, Hu G, Imwong M, Jaidee A, Russell B, et al. The transcriptome of Plasmodium vivax reveals divergence and diversity of transcriptional regulation in malaria parasites. Proc Natl Acad Sci USA. 2008;105: 16290–16295. doi:10.1073/pnas.0807404105

8. Caro F, Ahyong V, Betegon M, DeRisi JL. Genome-wide regulatory dynamics of translation in the Plasmodium falciparum asexual blood stages. eLife. 2014;3: 568. doi:10.7554/eLife.04106

9. Michel AM, Baranov PV. Ribosome profiling: a Hi-Def monitor for protein synthesis at the genome-wide scale. Wiley Interdiscip Rev RNA. 2013;4: 473–490. doi:10.1002/wrna.1172

10. Thiviyanathan V, Gorenstein DG. Aptamers and the next generation of diagnostic reagents. Stastna M, Van Eyk JE, editors. Proteomics Clin Appl. 2012;6: 563–573. doi:10.1002/prca.201200042

11. Oliphant AR, Brandl CJ, Struhl K. Defining the sequence specificity of DNA-binding proteins by selecting binding sites from random-sequence oligonucleotides: analysis of yeast GCN4 protein. Molecular and Cellular Biology. 1989;9: 2944–2949. doi:10.1128/mcb.9.7.2944

12. Tuerk C, Gold L. Systematic evolution of ligands by exponential enrichment: RNA ligands to bacteriophage T4 DNA polymerase. Science. 1990;249: 505–510.

13. Song H-O, Lee B, Bhusal RP, Park B, Yu K, Chong C-K, et al. Development of a novel fluorophore for real-time biomonitoring system. Carvalho LH, editor. PLoS ONE. 2012;7: e48459. doi:10.1371/journal.pone.0048459

14. Wang W-X, Cheung Y-W, Dirkzwager RM, Wong W-C, Tanner JA, Li H-W, et al. Specific and sensitive detection of Plasmodium falciparum lactate dehydrogenase by DNA-scaffolded silver nanoclusters combined with an aptamer. Analyst. 2017;142: 800–807. doi:10.1039/c6an02417c

15. Lee S, Manjunatha DH, Jeon W, Ban C. Cationic surfactant-based colorimetric detection of Plasmodium lactate dehydrogenase, a biomarker for malaria, using the specific DNA aptamer. Schallig HDFH, editor. PLoS ONE. 2014;9: e100847. doi:10.1371/journal.pone.0100847

16. Aurrecoechea C, Brestelli J, Brunk BP, Dommer J, Fischer S, Gajria B, et al. PlasmoDB: a functional genomic database for malaria parasites. Nucleic Acids Research. 2009;37: D539–43. doi:10.1093/nar/gkn814

17. Otto TD, Wilinski D, Assefa S, Keane TM, Sarry LR, Böhme U, et al. New insights into the blood-stage transcriptome of Plasmodium falciparum using RNA-Seq. Molecular Microbiology. 2010;76: 12–24. doi:10.1111/j.1365-2958.2009.07026.x

18. Gelinas AD, Davies DR, Janjic N. Embracing proteins: structural themes in aptamer-protein complexes. Curr Opin Struct Biol. 2016;36: 122–132. doi:10.1016/j.sbi.2016.01.009

19. Štros M, Launholt D, Grasser KD. The HMG-box: a versatile protein domain occurring in a wide variety of DNA-binding proteins. Cell Mol Life Sci. 2007;64: 2590–2606. doi:10.1007/s00018-007-7162-3

20. Sievers F, Wilm A, Dineen D, Gibson TJ, Karplus K, Li W, et al. Fast, scalable generation of high-quality protein multiple sequence alignments using Clustal Omega. Mol Syst Biol. 2011;7: 539–539. doi:10.1038/msb.2011.75

21. Milon P, Konevega AL, Peske F, Fabbretti A, Gualerzi CO, Rodnina MV. Transient kinetics, fluorescence, and FRET in studies of initiation of translation in bacteria. Meth Enzymol. 2007;430: 1–30. doi:10.1016/S0076-6879(07)30001-3

22. Lim TS, Schütze T, Lehrach H, Glökler J, Konthur Z. Diversity visualization by endonuclease: a rapid assay to monitor diverse nucleotide libraries. Anal Biochem. 2011;411: 16–21. doi:10.1016/j.ab.2010.12.024

23. Alam KK, Chang JL, Burke DH. FASTAptamer: A Bioinformatic Toolkit for High-throughput Sequence Analysis of Combinatorial Selections. Mol Ther Nucleic Acids. 2015;4: e230. doi:10.1038/mtna.2015.4

24. Bailey TL, Boden M, Buske FA, Frith M, Grant CE, Clementi L, et al. MEME SUITE: tools for motif discovery and searching. Nucleic Acids Research. 2009;37: W202–8. doi:10.1093/nar/gkp335

25. Zuker M. Mfold web server for nucleic acid folding and hybridization prediction. Nucleic Acids Research. 2003;31: 3406–3415. doi:10.1093/nar/gkg595

26. Jeddi I, Saiz L. Three-dimensional modeling of single stranded DNA hairpins for aptamer-based biosensors. Sci Rep. 2017;7: 1178. doi:10.1038/s41598-017-01348-5

27. Popenda M, Szachniuk M, Antczak M, Purzycka KJ, Lukasiak P, Bartol N, et al. Automated 3D structure composition for large RNAs. Nucleic Acids Research. 2012;40: e112–e112. doi:10.1093/nar/gks339

28. Humphrey W, Dalke A, Schulten K. VMD: visual molecular dynamics. J Mol Graph. 1996;14: 33–8-27–8.

29. MacKerell AD, Banavali N, Foloppe N. Development and current status of the CHARMM force field for nucleic acids. Biopolymers. 2000;56: 257–265. doi:10.1002/1097-0282(2000)56:4<257::AID-BIP10029>3.0.CO;2-W

30. Phillips JC, Braun R, Wang W, Gumbart J, Tajkhorshid E, Villa E, et al. Scalable molecular dynamics with NAMD. J Comput Chem. 2005;26: 1781–1802. doi:10.1002/jcc.20289

31. Waterhouse A, Bertoni M, Bienert S, Studer G, Tauriello G, Gumienny R, et al. SWISS-MODEL: homology modelling of protein structures and complexes. Nucleic Acids Research. 2018;46: W296–W303. doi:10.1093/nar/gky427

32. Pierce BG, Wiehe K, Hwang H, Kim B-H, Vreven T, Weng Z. ZDOCK server: interactive docking prediction of protein-protein complexes and symmetric multimers. Bioinformatics. 2014;30: 1771–1773. doi:10.1093/bioinformatics/btu097

33. Chulluncuy R, Espiche C, Nakamoto JA, Fabbretti A, Milon P. Conformational Response of 30S-bound IF3 to A-Site Binders Streptomycin and Kanamycin. Antibiotics (Basel). Multidisciplinary Digital Publishing Institute; 2016;5: 38. doi:10.3390/antibiotics5040038

34. Milon P, Konevega AL, Gualerzi CO, Rodnina MV. Kinetic checkpoint at a late step in translation initiation. Molecular Cell. 2008;30: 712–720. doi:10.1016/j.molcel.2008.04.014

35. Ingolia NT, Ghaemmaghami S, Newman JRS, Weissman JS. Genome-wide analysis in vivo of translation with nucleotide resolution using ribosome profiling. Science. American Association for the Advancement of Science; 2009;324: 218–223. doi:10.1126/science.1168978

36. Kim H, Certa U, Döbeli H, Jakob P, Hol WG. Crystal structure of fructose-1,6-bisphosphate aldolase from the human malaria parasite Plasmodium falciparum. Biochemistry. 1998;37: 4388–4396. doi:10.1021/bi972233h

37. Ellington AD, Szostak JW. In vitro selection of RNA molecules that bind specific ligands. Nature. 1990;346: 818–822. doi:10.1038/346818a0

38. Stoltenburg R, Reinemann C, Strehlitz B. FluMag-SELEX as an advantageous method for DNA aptamer selection. Anal Bioanal Chem. 2005;383: 83–91. doi:10.1007/s00216-005-3388-9

39. Schneider CA, Rasband WS, Eliceiri KW. NIH Image to ImageJ: 25 years of image analysis. Nature Methods. 2012;9: 671–675. doi:10.1038/nmeth.2089

40. Duhr S, Braun D. Why molecules move along a temperature gradient. Proceedings of the National Academy of Sciences. 2006;103: 19678–19682. doi:10.1073/pnas.0603873103

41. Dhont JKG, Wiegand S, Duhr S, Braun D. Thermodiffusion of charged colloids: single-particle diffusion. Langmuir. 2007;23: 1674–1683. doi:10.1021/la062184m

42. Breitsprecher D, Schlinck N, Witte D, Duhr S, Baaske P, Schubert T. Aptamer Binding Studies Using MicroScale Thermophoresis. Methods Mol Biol. New York, NY: Springer New York; 2016;1380: 99–111. doi:10.1007/978-1-4939-3197-2_8

43. Hianik T, Ostatná V, Sonlajtnerova M, Grman I. Influence of ionic strength, pH and aptamer configuration for binding affinity to thrombin. Bioelectrochemistry. 2007;70: 127–133. doi:10.1016/j.bioelechem.2006.03.012

44. Bell D, Peeling RW, WHO-Regional Office for the Western Pacific/TDR. Evaluation of rapid diagnostic tests: malaria. Nature reviews. Microbiology. 2006. pp. S34–8. doi:10.1038/nrmicro1524

45. Kaur H, Sehgal R, Kumar A, Sehgal A, Bansal D, Sultan AA. Screening and identification of potential novel biomarker for diagnosis of complicated Plasmodium vivax malaria. Journal of Translational Medicine. BioMed Central; 2018;16: 42. doi:10.1186/s12967-018-1646-9

46. Meerstein-Kessel L, van der Lee R, Stone W, Lanke K, Baker DA, Alano P, et al. Probabilistic data integration identifies reliable gametocyte-specific proteins and transcripts in malaria parasites. Sci Rep. Nature Publishing Group; 2018;8: 410. doi:10.1038/s41598-017-18840-7

47. Frith K-A, Fogel R, Goldring JPD, Krause RGE, Khati M, Hoppe H, et al. Towards development of aptamers that specifically bind to lactate dehydrogenase of Plasmodium falciparum through epitopic targeting. Malar J. 2018;17: 191. doi:10.1186/s12936-018-2336-z

48. Rousu J, Agranoff DD, Sodeinde O, Shawe-Taylor J, Fernandez-Reyes D. Biomarker discovery by sparse canonical correlation analysis of complex clinical phenotypes of tuberculosis and malaria. Price ND, editor. PLoS Comput Biol. 2013;9: e1003018. doi:10.1371/journal.pcbi.1003018

49. Kumar K, Singal A, Rizvi MMA, Chauhan VS. High mobility group box (HMGB) proteins of Plasmodium falciparum: DNA binding proteins with pro-inflammatory activity. Parasitol Int. 2008;57: 150–157. doi:10.1016/j.parint.2007.11.005

50. Klune JR, Dhupar R, Cardinal J, Billiar TR, Tsung A. HMGB1: endogenous danger signaling. Mol Med. 2008;14: 476–484. doi:10.2119/2008-00034.Klune

51. Briquet S, Lawson-Hogban N, Boisson B, Soares MP, Péronet R, Smith L, et al. Disruption of Parasite hmgb2 Gene Attenuates Plasmodium berghei ANKA Pathogenicity. Infection and Immunity. 2015;83: 2771–2784. doi:10.1128/IAI.03129-14

52. Nimjee SM, White RR, Becker RC, Sullenger BA. Aptamers as Therapeutics. Annu Rev Pharmacol Toxicol. 2017;57: 61–79. doi:10.1146/annurev-pharmtox-010716-104558

53. Cheung Y-W, Dirkzwager RM, Wong W-C, Cardoso J, D’Arc Neves Costa J, Tanner JA. Aptamer-mediated Plasmodium-specific diagnosis of malaria. Biochimie. 2018;145: 131–136. doi:10.1016/j.biochi.2017.10.017

54. Miranda GF, Feng L, Shiu SC-C, Dirkzwager RM, Cheung Y-W, Tanner JA, et al. Aptamer-Based Electrochemical Biosensor for Highly Sensitive and Selective Malaria Detection with Adjustable Dynamic Response Range and Reusability. Sensors & Actuators: B Chemical. Elsevier B.V; 2017;: 1–23. doi:10.1016/j.snb.2017.07.117

55. Hamaguchi N, Ellington A, Stanton M. Aptamer beacons for the direct detection of proteins. Anal Biochem. 2001;294: 126–131. doi:10.1006/abio.2001.5169

56. Gawande BN, Rohloff JC, Carter JD, Carlowitz von I, Zhang C, Schneider DJ, et al. Selection of DNA aptamers with two modified bases. Proc Natl Acad Sci USA. 2017;114: 2898–2903. doi:10.1073/pnas.1615475114

57. Abeydeera ND, Egli M, Cox N, Mercier K, Conde JN, Pallan PS, et al. Evoking picomolar binding in RNA by a single phosphorodithioate linkage. Nucleic Acids Research. 2016;44: 8052–8064. doi:10.1093/nar/gkw725

